# Dual strategies in ant social immunity: Chemical diversity and pathogen specificity in antimicrobial defense

**DOI:** 10.1101/2025.07.24.666674

**Authors:** Mary K. Chon, Darmon Kahvazadeh, Clint A. Penick

**Affiliations:** Department of Ecology, Evolution, & Organismal Biology, Kennesaw State University, Kennesaw, GA, USA; Department of Entomology and Plant Pathology, Auburn University, Auburn, AL, USA

**Keywords:** social immunity, antimicrobials, antibiotics, *Candida auris*, social insects, ants

## Abstract

The evolution of sociality provides many benefits but also increases the risk of disease transmission. In response, social insects have evolved social immune defenses, which includes the production and communal use of antimicrobial compounds. The origins of sociality in insects are closely associated with the development of exocrine glands that secrete these antimicrobials. However, it is unclear how social insects prevent pathogens from evolving resistance to these compounds given their frequent use. We tested whether ants reduce this risk by producing chemically diverse and pathogen-specific antimicrobial compounds. Using solvents of varying polarity, we extracted antimicrobial compounds from six ant species and tested their inhibitory effects on Gram-positive bacteria, Gram-negative bacteria, and a fungal pathogen. Our results show that ants produce multiple classes of antimicrobial compounds with distinct chemical properties and targeted activity, though the strength and specificity of inhibition vary across species. Notably, five of the six ant species tested inhibited *Candida auris*, an emerging fungal pathogen of critical concern in human medicine due to its multidrug resistance. These findings suggest that social immunity in ants is reinforced by both chemical diversity and pathogen specificity, offering an evolutionary strategy that may help prevent antimicrobial resistance in dense, socially complex environments.

## Introduction

Social organisms face stronger disease pressures due to their dense living conditions and high relatedness among group members (Pie, Rosengaus, & Traniello, 2004; Böhm *et al.*, 2008; Kappeler, Cremer, & Nunn, 2015). In humans, for example, dense living conditions in cities create hot spots for the rapid spread of disease (Baker *et al.*, 2022). Similarly, social insects (ants, bees, wasps, and termites) live in densely populated societies where disease transmission risk is high (Pie *et al.*, 2004; Meunier, 2015). To counteract these risks, social organisms have developed a range of disease defenses that collectively function as social immunity—a group-level defense system particularly well-developed in social insects (Cremer, Armitage, & Schmid-Hempel, 2007; Cremer, Pull, & Fürst, 2018). These defenses include hygienic behaviors as well as the secretion of antimicrobial compounds from specialized glands, which are applied to nestmates, brood, and nest surfaces to prevent infection (Beattie *et al.*, 1986; Stow *et al.*, 2007; Hoggard *et al.*, 2011; Tragust, 2016). While the rise of antimicrobial resistance poses a growing challenge in medicine, social insects appear to have used chemical defenses effectively for millions of years. Yet it remains unclear how these long-standing antimicrobial strategies have avoided driving resistance in the pathogens they target.

The development of antimicrobials has been identified as a key adaptation in social insects to facilitate group living (Meunier, 2015). Theory about social evolution has predicted that social species will invest more heavily in antimicrobials than solitary species. This has been supported by past research comparing antimicrobial investment among social and solitary wasps and bees (Stow *et al.*, 2007; Hoggard *et al.*, 2011). Among ants—all of which are social—the majority of species produce potent antimicrobials, though investment varies (Penick *et al.*, 2018). Ants possess multiple antimicrobial-producing glands, including the metapleural (Maschwitz, Koob, & Schildknecht, 1970; Poulsen *et al.*, 2002; Fernández-Marín *et al.*, 2006; Yek *et al.*, 2012), poison (Jouvenaz, Blum, & MacConnell, 1972; Blum, 1992; Tragust *et al.*, 2013), mandibular (Brough, 1983), and Dufour’s (Quinet *et al.*, 2012) glands. The diversity of these glands underscores the importance of antimicrobial defense in ant evolution and their interactions with bacterial and fungal pathogens (Vander Meer, 2012).

While broad-spectrum antimicrobials have been widely studied in social insects, it remains unclear how they prevent pathogens from evolving resistance. We propose two potential strategies: (1) producing multiple classes of antimicrobial compounds with distinct chemical properties, making it harder for pathogens to adapt, and (2) employing pathogen-specific antimicrobials that selectively target different microbial threats. These approaches parallel human antimicrobial strategies, where drug cocktails (e.g., combination antibiotic therapy) and targeted treatments (e.g., antifungal or antiviral drugs) help slow resistance evolution (Chait, Craney, & Kishony, 2007; Maxson & Mitchell, 2016; Singh & Yeh, 2017). Testing these strategies in ants offers a model for understanding how long-term use of antimicrobials can be sustained without promoting resistance.

Evidence suggests social insects may employ both strategies. The metapleural glands of ants produce a variety of antimicrobial compounds, including carboxylic acids, alkylphenols, and antifungals (Beattie *et al.*, 1986; Yek *et al.*, 2012; Tragust, 2016). Their venom also contains antimicrobial compounds, such as alkaloids, decahydroquinolines, and antimicrobial peptides (Moreau, 2013; Das *et al.*, 2014; Tragust, 2016; Yacoub *et al.*, 2020). Additionally, some insect antimicrobials also show specificity against particular pathogens. For example, termites enhance antimicrobial activity after exposure to drug-resistant bacteria such as methicillin-resistant *Staphylococcus aureus* (MRSA) (Zeng, Hu, & Suh, 2016). Clarifying whether ants deploy both broad-spectrum and pathogen-specific antimicrobials is essential to understanding how they limit pathogen resistance. If ants rely on multiple compound classes with distinct chemical properties and selectively target different microbial threats, this dual strategy could represent a robust evolutionary solution to the challenge of antimicrobial resistance.

Here, we test whether ants employ these two strategies by analyzing antimicrobial extracts from six ant species that vary in life history and antimicrobial production (**Table 1**). We extracted putative antimicrobials using solvents of varying polarity and tested their activity against three major pathogen groups (Gram-positive bacteria, Gram-negative bacteria, and fungi). Using solvents that differ in polarity allows us to extract a broad range of compounds with diverse chemical properties, providing a more comprehensive assessment of whether ants produce multiple classes of antimicrobials. If ants employ this strategy, we predict that antimicrobial extracts obtained using both polar and non-polar solvents will inhibit microbial growth. Additionally, if ants use antimicrobials that vary in microbe specificity, we expect that some species will produce antimicrobials that effectively inhibit certain microbial groups but not others.

**Table 1.** Traits of focal ant species and their putative antimicrobials (from focal species or their close relatives).

| Ant species | Colony size* | Foraging niche<sup>†</sup> | Nesting niche<sup>†</sup> | Diet<sup>‡</sup> | Putative antimicrobials<sup>‡</sup> |
| --- | --- | --- | --- | --- | --- |
| *Crematogaster ashmeadi* Mayr | 10,000 | Arboreal | Carton | Omnivorous | Phenols and carboxylic acids |
| *Solenopsis invicta* Buren | 220,000 | Epigaeic | Ground | Omnivorous | Piperidine Alkaloids (e.g., solenopsin) |
| *Prenolepis imparis* (Say) | 3,370 | Epigaeic | Ground | Omnivorous | Formic acid |
| *Linepithema humile* (Mayr) | 150,000 | Epigaeic | Ground | Omnivorous | Iridoids (e.g., dolichodial, iridomyrmecin) |
| *Dorymyrmex bureni* (Trager) | 1,000 | Epigaeic | Ground | Omnivorous | Non-alkaloidal amines (e.g., actadine) |
| *Brachyponera chinensis* (Emery) | 1,000 | Epigaeic | Leaf Litter | Predator | Unknown |

\*References for colony size from Penick et al. (2018) <sup>†</sup>Categorized based on definitions from Sosiak and Barden (2021) <sup>‡</sup>References for antimicrobials include Yan et al. (2017), Wang et al. (2015), Marlier et al. (2004), Bai et al. (2024), Kesäniemi et al. (2019), Dioguardi et al. (2024), and Touchard et al. (2016)

## Methods

### Ant collection

To evaluate antimicrobial inhibition, we collected workers of six ant species from independent nests in Alabama, Florida, and Georgia (USA) from February 2023 to June 2024 (see **Table S1** for detailed collection information). These six ant species included representatives from all four major ant subfamilies (Ponerinae, Dolichoderinae, Formicinae, and Myrmecinae) and included species that were previously shown to vary in their antimicrobial defenses against *Staphylococcus epidermidis* (Penick *et al.*, 2018). At least 60 workers were collected from each independent nest for antimicrobial trials, and we excluded young, callow individuals. We ran 11-43 independent nests per species, 122 nests total (see **Table S2** for sample sizes for each test independently). Voucher specimens from each nest were reserved in 70% ethanol for identification using the keys from Mississippi State University (MacGown, 2020).

### Antimicrobial extraction

We soaked groups of 15 ants from each colony for 24 hours in one of four solvents selected based on their polarity (from most to least polar): 95% ethanol (Koptec, PA, USA), isopropanol, dichloromethane, and hexane (all from VWR Chemicals, PA, USA). We filtered extracts through sterile 0.2μM polyethersulfone filters into newly labeled 1.5mL microtubes that we then dried using a Eppendorf^®^ Vacufuge Plus concentrating centrifuge over 6-12hrs. The dried extracts were resuspended in 400 μL of sterile Luria-Bertani (LB) broth, vortexed, and immediately used for antimicrobial assays.

### Antimicrobial assay

Using methods modified from Penick *et al.* (Penick *et al.*, 2018), we quantified growth inhibition of ant extracts against three test microbes: *Staphylococcus epidermidis* strain 12228 (Gram-positive bacterium)*, Escherichia coli* strain ATCC 25922 (Gram-negative bacterium), and *Candida auris* strain AR.0381 (fungal pathogen). These strains were selected to represent a range of potential microbes ants may encounter in their natural environments*. S. epidermidis* and *E. coli* are common model organisms for Gram-positive and Gram-negative bacteria, respectively, and are frequently used to assess broad-spectrum antimicrobial activity. *C. auris* was included as a representative fungal pathogen that more closely approximates entomopathogenic fungi, although most ant-infecting fungi are filamentous and not compatible with a plate reader assay (Baron *et al.*, 2019).

For each test microbe, we added 100 µL of ant extract (3.75-ant equivalent) into separate wells of a 96-well plate containing 100 µL of Luria-Bertani (LB) broth with 100 colony-forming units (CFUs) of the respective microbe (**Fig. S1**). CFU counts were verified by plating serial 1:100 dilutions of each pathogen onto nine plates and comparing counted CFUs to optical density (OD) measurements. Since each extract was tested against all three microbes, a total of 300 µL of ant extract was used per trial. To ensure sterility, an additional 100 µL of each ant extract was added to wells containing 100 µL of LB without microbes. Three control wells per plate contained 200 µL of LB with 100 CFUs of each test microbe to serve as positive growth controls.

Microbial growth was recorded hourly over 24 hours using a microplate reader (Molecular Devices, SpectraMax^®^ 340PC 384, CA, USA) after shaking to resuspend cultures. Inhibition was assessed as an all-or-none response: if the change in OD from the initial to peak reading over 24 hours was less than 10% of the OD change observed in positive control wells, the sample was considered completely inhibitory. Inhibition was calculated using the equation:

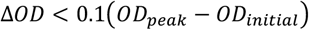

where OD*_initial_* is the OD of positive control wells at time zero, and OD*_peak_* is the highest, non-outlier OD value recorded. Samples were excluded from inhibition analysis if microbial growth occurred in sterility control wells or if no growth was observed in positive control wells. We chose this binary scoring method because the assay is highly sensitive and optimized for small volumes of ant extract. Using a conservative threshold to define inhibition reduces variability across trials and allows for consistent comparisons among species. Previous studies have validated this method as effective for detecting species-level differences in antimicrobial investment (Penick *et al.*, 2018; Halawani *et al.*, 2020). We standardized extract concentration to 3.75 ant equivalents per trial, based on prior work indicating that this amount yields the clearest interspecific differences (Penick *et al.*, 2018).

### Statistical analyses

All statistical analyses were performed using R v4.4.2 (R Core Team, 2025). To test whether antimicrobial activity differed among ants and varied by solvent and test microbe, we used a generalized linear mixed model (GLMM) with a binomial distribution and logit link function using the ‘lme4’ package (Bates *et al.*, 2015). Subfamily was included as a random effect to account for phylogenetic non-independence among ant species. Post-hoc pairwise comparisons were conducted using estimated marginal means (EMMs) with the ‘emmeans’ package (Lenth, 2025). Comparisons were made within each species for both solvent and test microbe treatments. P-values were corrected for multiple comparisons using the Benjamini-Hochberg (FDR) method.

## Results

### Antimicrobial activity

Ant species differed significantly in their ability to inhibit microbial growth, with inhibition patterns varying by both solvent type and test microbe (**Table 2**). Most species showed inhibition with both polar and non-polar solvents, supporting our first prediction that ants produce multiple classes of antimicrobial compounds with distinct chemical properties (**Table S3**; **Fig. 1)**. Significant pairwise contrasts (**Table S4**; **Fig. 2A–F**) confirmed stronger inhibition with polar solvents (ethanol and isopropanol), particularly in *Solenopsis invicta, Crematogaster ashmeadi*, and *Brachyponera chinensis*. In contrast, *Dorymyrmex bureni* showed greater inhibition of *S. epidermidis* and *E. coli* with non-polar solvents (dichloromethane and hexane), although these effects were only marginally significant (**Table S4**).

**Table 2.** Generalized linear mixed model (GLMM) results.

| Comparison | Chi<sup>2</sup> | df | p-value |
| --- | --- | --- | --- |
| Ant species | 40.22 | 5 | 0.0061** |
| Solvent | 23.33 | 3 | 0.0067** |
| Test microbe | 35.085 | 2 | < 0.0001*** |
| Ant species*solvent | 131.20 | 15 | < 0.0001*** |
| Ant species*test microbe | 34.12 | 10 | 0.0116** |

**Figure 1.**
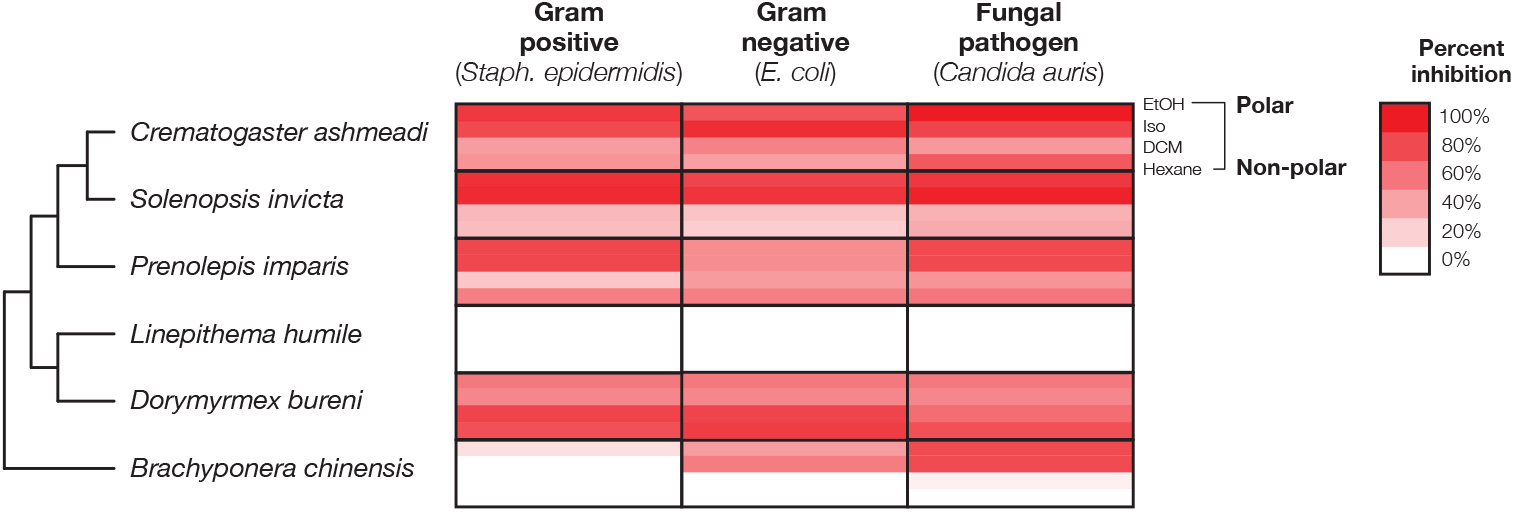
Antimicrobial activity of ant extracts. Colors indicate percent of trials in which growth was fully inhibited of each of three microbes using four extracts that vary in polarity (from most to least polar: ethanol [EtOH], isopropanol [Iso], dichloromethane [DCM], and hexane [Hex]). Percent inhibition is indicated using a color gradient, with red indicating inhibition observed in 100% of trials, while white indicates no inhibition; inhibition shows the percentage of independent colonies per test that exhibited inhibition. The phylogeny showing relatedness among ants’ species is based on Blanchard and Moreau (Blanchard & Moreau, 2017).

**Figure 2.**
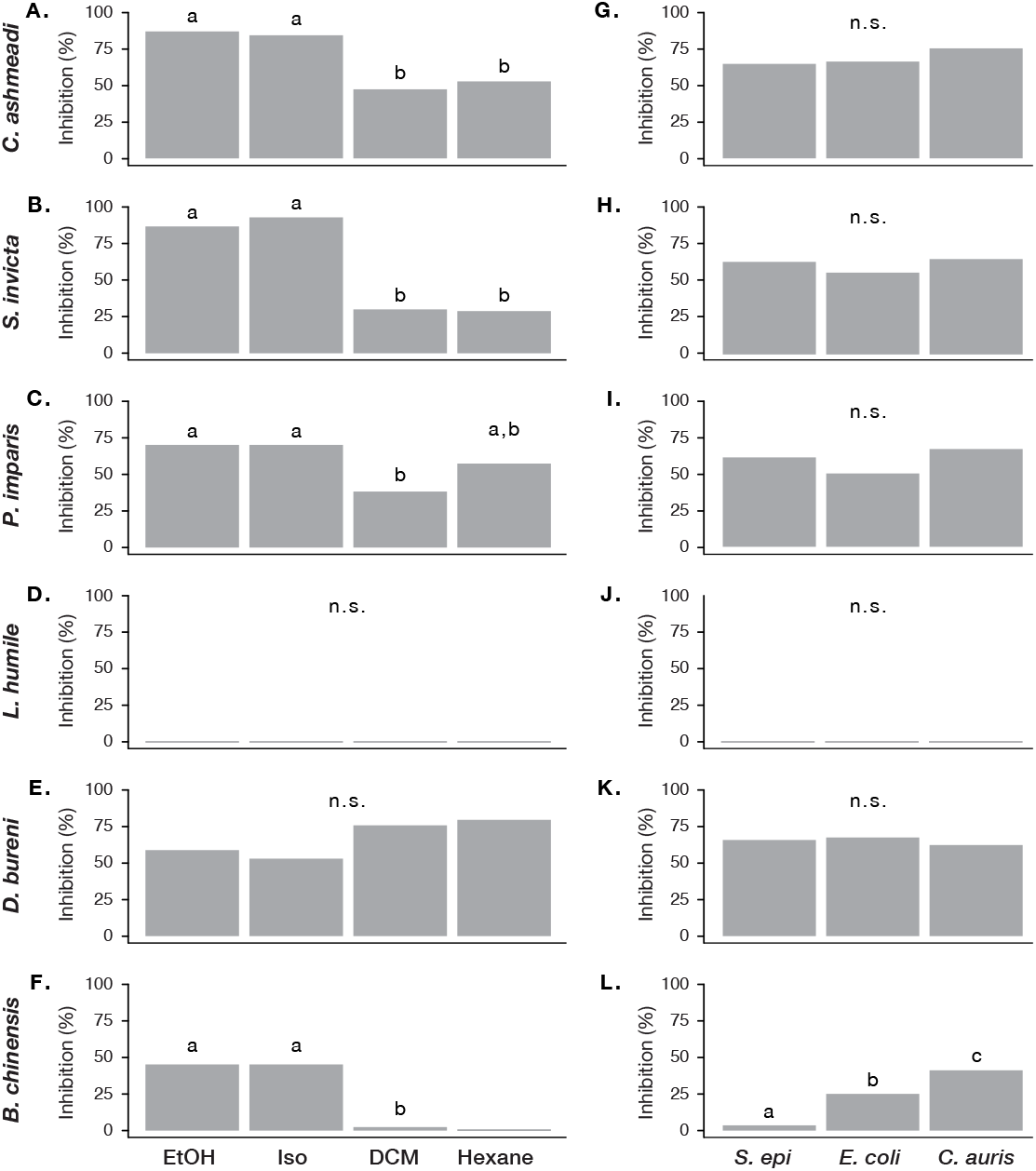
Variation in antimicrobial activity across solvents (**A–F**) and test microbes (**G–L**) for each ant species. Bars show the percentage of trials exhibiting microbial inhibition. Different letters denote statistically significant pairwise differences between groups (p < 0.05).

Four of the six focal species (*S. invicta, C. ashmeadi, D. bureni*, and *P. imparis*) demonstrated broad-spectrum antimicrobial activity, inhibiting all three test microbes in 45–75% of trials when combining results across all solvents (**Table S5**; **Fig. 2G–L**). *L. humile* exhibited no inhibition under any treatment. *B. chinensis* showed mixed activity, significantly inhibiting *E. coli* and *C. auris* but exhibiting reduced inhibition against *S. epidermidis* (**Fig. 2L**). These findings partially support our second prediction: some species (e.g., *B. chinensis*) may produce pathogen-specific antimicrobials, while others (e.g., *S. invicta, C. ashmeadi*) invest in broad-spectrum defenses (**Table 3**). Additionally, *D. bureni* and *P. imparis* showed evidence of pathogen-specific inhibition that varied by solvent (**Fig. 1**), suggesting these species may also produce chemically distinct antimicrobials that target different microbial threats. The complete absence of inhibition by *L. humile* suggests it either lacks external antimicrobial defenses or employs an alternative strategy.

**Table 3.** Summary of inhibition results across species.

| Species | Broad-spectrum | Solvent effect | Microbe effect |
| --- | --- | --- | --- |
| *Crematogaster ashmeadi* | Yes | Yes | No |
| *Solenopsis invicta* | Yes | Yes | No |
| *Prenolepis imparis* | Partial | Yes | No |
| *Linepithema humile* | None | No | No |
| *Dorymyrmex bureni* | Yes | No | No |
| *Brachyponera chinensis* | Partial | Yes | Yes |

## Discussion

Antimicrobial resistance poses a significant challenge in both human medicine and natural systems (Woolhouse *et al.*, 2015), yet social insects have seemingly employed antimicrobial defenses for millions of years without widespread pathogen resistance. Our study found support for two potential strategies ants may use to prevent antimicrobial resistance. First, we found that polar and non-polar extracts exhibited antimicrbial activity, which indicates that ants produce multiple classes of antimicrobial compounds with distinct chemical properties. Second, we found that ants produced antimicrbials active against specific classes of potential pathogens, which suggests that ants invest in pathogen-specific antimicrobials that selectively target different microbial threats. These approaches parallel human antimicrobial strategies, where combination therapies and targeted treatments help slow resistance evolution (Chait *et al.*, 2007; Maxson & Mitchell, 2016; Singh & Yeh, 2017).

Our results provide strong evidence that ants employ chemically diverse antimicrobial compounds, supporting our first hypothesis. Extracts from polar solvents (ethanol and isopropanol) exhibited the strongest antimicrobial activity, consistent with previous findings that show ants invest in water-soluble antimicrobial compounds, such as antimicrobial peptides, venom alkaloids, or formic acid (Jouvenaz *et al.*, 1972; Téné *et al.*, 2016; Brütsch *et al.*, 2017; Penick *et al.*, 2018). The widespread use of polar antimicrobials in insects and plants is well documented, as these compounds readily dissolve in water and interact with microbial cell walls (Fan, Ngo, & Wu, 2018). However, non-polar extracts (dichloromethane and hexane) also exhibited antimicrobial activity in several of the species we tested, suggesting that lipid-soluble compounds contribute to antimicrobial defense. Non-polar antimicrobial compounds may be particularly relevant for insects, whose exoskeletons are coated in lipid-rich cuticular hydrocarbons that serve as barriers to water loss and may play roles in communication and defense (Koidsumi, 1957; Blomquist & Bagnères, 2010). The presence of both polar and non-polar antimicrobial compounds in ants suggests a sophisticated chemical strategy, potentially reducing the likelihood of pathogen adaptation.

Our results also support the second hypothesis that ants produce antimicrobials that selectively inhibit different classes of microbes. Three ant species (*S. invicta, C. ashmeadi*, and *D. bureni*) exhibited broad-spectrum antimicrobial activity, inhibiting all three microbial groups (Gram-positive bacteria, Gram-negative bacteria, and fungi). However, *B. chinensis* inhibited *E. coli* and *C. auris* but not *S. epidermidis*, while *P. imparis* inhibited *C. auris* and *S. epidermidis* but had reduced inhibition against *E. coli*. These findings suggest that some ants selectively invest in antimicrobials that target specific microbial threats, a strategy analogous to targeted antimicrobial therapies in human medicine (Maxson & Mitchell, 2016). It remains unclear whether ants actively regulate their antimicrobial production in response to environmental cues or whether these strategies are fixed. In termites, exposure to drug-resistant bacteria has been shown to enhance antimicrobial activity (Zeng *et al.*, 2016), but a similar study in ants found no evidence for inducible antimicrobial defenses (Halawani *et al.*, 2020). Our findings suggest that at least some ants may constitutively produce pathogen-specific antimicrobials rather than modulating their production in response to microbial threats. Future research should investigate whether ants can alter antimicrobial production based on infection risk and whether specific glands contribute to different antimicrobial strategies.

Beyond our central hypotheses, we also found that five of the six ant species tested produced compounds that fully inhibited *Candida auris*, a fungal pathogen of urgent concern in human medicine due to its multidrug resistance and rising prevalence in healthcare settings (Chowdhary, Sharma, & Meis, 2017). This finding suggests that ants, which likely experience strong selection from fungal pathogens in their environments, may serve as an underexplored source of antifungal compounds. Given the growing demand for novel antifungal agents, particularly those effective against *C. auris* (Lockhart, 2019), ant-derived chemicals could offer promising leads for future pharmaceutical development.

Notably, *L. humile* exhibited no detectable antimicrobial activity against any of the tested microbes. Given that many ant species are known to produce potent antimicrobials, this finding is unexpected—especially for a globally invasive species (Silverman & Brightwell, 2008). One possibility is that it does produce antimicrobials, but they were not captured by our assay. Some insects only express antimicrobials after pathogen exposure (Zeng *et al.*, 2016), while others produce volatile compounds that may evade detection in plate-based assays (Wang *et al.*, 2015). Alternatively, *L. humile* may rely on nonchemical disease defenses. Social insects can limit pathogen spread through behaviors such as grooming, corpse removal, and restructuring social networks (Cremer *et al.*, 2018), or by emphasizing internal immune responses over external antimicrobial production. Interestingly, species that lack external antimicrobials have sometimes shown higher survival following microbial exposure than those that invest heavily in chemical defenses (Halawani *et al.*, 2020). Some ants also harbor beneficial microbial symbionts that produce antibiotics, as seen in fungus-growing species (Little *et al.*, 2006; Mendes *et al.*, 2013; Henrik, Boomsma, & Tunlid, 2014), though the extent of such symbioses in other ants remains poorly understood. Future research should explore whether *L. humile* compensates for its apparent lack of antimicrobial investment through behavioral defenses or microbial partnerships.

Our findings contribute to broader understanding of social immunity by showing that ants not only produce diverse antimicrobials but also deploy them in species-specific ways that likely reflect different colony-level strategies for managing microbial threats. The presence of both broad-spectrum and pathogen-specific antimicrobials across species suggests variation in how social insects invest in external defenses—a key component of social immunity. These results imply that selection has favored flexibility in antimicrobial deployment, allowing colonies to adapt their chemical defense portfolios in response to ecological context. As such, antimicrobial diversity may represent a chemical form of social immunity shaped by evolutionary pressures associated with group living.

## Acknowledgements

For help with ant collection, rearing, and antimicrobial assays, we thank Victoria Garcia-Belman, Cristian Rodriguez, Dawson Hicks, Brian Dodson, Alyssa Dance, Tyler Thompson, Jayce Easterwood, and Adalynn Chon. We thank Thomas McElroy, Christopher Cornelison, and Andrew Haddow for helpful input on an earlier version of this manuscript. This project is based on research that was partially supported by the Alabama Agricultural Experiment Station with funding from the Hatch Act capacity funding program (Accession Number 7008193) from the USDA National Institute of Food and Agriculture as well as the Agriculture Research Enhancement and Seed Funding Program awarded to CP.

## Data availability statement

All data and analysis code supporting the findings of this study are available in the Supplementary Material associated with this article.

## References

Bai W, Chen B, Chen H, Nie L, Liang M, Xu Y, Lu Y & Wang L. 2024. Antimicrobial Activity of Compounds Isolated from the Nest Material of Crematogaster rogenhoferi (Mayr) (Hymenoptera: Formicidae). Insects 15: 1019.

Baker RE, Mahmud AS, Miller IF, Rajeev M, Rasambainarivo F, Rice BL, Takahashi S, Tatem AJ, Wagner CE, Wang LF, Wesolowski A & Metcalf CJE. 2022. Infectious disease in an era of global change. Nature Reviews Microbiology 20: 193–205.

Baron NC, Rigobelo EC, Zied DC, Baron NC, Rigobelo EC & Zied DC. 2019. Filamentous fungi in biological control: Current status and future perspectives. Chilean journal of agricultural research 79: 307–315.

Bates D, Maechler M, Bolker B, Walker S, Christensen RHB, Singmann H, Dai B, Grothendieck G, Green P & Bolker B. 2015. Package ‘lme4’.: 1.1-35.5.

Beattie AJ, Turnbull CL, Hough T & Knox RB. 1986. Antibiotic production: A possible function for the metapleural glands of ants (Hymenoptera: Formicidae). Annals of the Entomological Society of America 79: 448–450.

Blanchard BD & Moreau CS. 2017. Defensive traits exhibit an evolutionary trade-off and drive diversification in ants. Evolution 71: 315–328.

Blomquist GJ & Bagnères AG. 2010. Insect Hydrocarbons: Biology, Biochemistry, and Chemical Ecology. Cambridge, UK: Cambridge University Press.

Blum MS. 1992. Ant venoms: Chemical and pharmacological properties. Journal of Toxicology: Toxin Reviews 11: 115–164.

Böhm M, Palphramand KL, Newton-Cross G, Hutchings MR & White PCL. 2008. Dynamic interactions among badgers: Implications for sociality and disease transmission. Journal of Animal Ecology 77: 735– 745.

Brough EJ. 1983. The antimicrobial activity of the mandibular gland secretion of a formicine ant, Calomyrmex sp. (Hymenoptera: Formicidae). Journal of Invertebrate Pathology 42: 306–311.

Brütsch T, Jaffuel G, Vallat A, Turlings TCJ & Chapuisat M. 2017. Wood ants produce a potent antimicrobial agent by applying formic acid on tree-collected resin. Ecology and Evolution 7: 2249–2254.

Chait R, Craney A & Kishony R. 2007. Antibiotic interactions that select against resistance. Nature 446: 668–671.

Chowdhary A, Sharma C & Meis JF. 2017. Candida auris: A rapidly emerging cause of hospital-acquired multidrug-resistant fungal infections globally. PLOS Pathogens 13: e1006290.

Cremer S, Armitage SA & Schmid-Hempel P. 2007. Social immunity. Current biology 17: R693–R702.

Cremer S, Pull CD & Fürst MA. 2018. Social immunity: Emergence and evolution of colony-level disease protection. Annual Review of Entomology 63: 105–123.

Das P, Dileepkumar R, Anaswara Krishnan S, Nair AS, Dhar PK & Oommen OV. 2014. Decahydroquinolines from the venom of a formicinae ant, Oecophylla smaragdina. Toxicon 92: 50–53.

Dioguardi M, Cantore S, Sovereto D, Sanesi L, Martella A, Almasri L, Musella G, Lo Muzio L & Ballini A. 2024. Therapeutic Potential of Solenopsis invicta Venom: A Scoping Review of Its Bioactive Molecules, Biological Aspects, and Health Applications. Biomolecules 14: 1499.

Fan X, Ngo H & Wu C. 2018. Natural and bio-based antimicrobials: A review. In: Natural and Bio-Based Antimicrobials for Food Applications. American Chemical Society, 1–24.

Fernández-Marín H, Zimmerman JK, Rehner SA & Wcislo WT. 2006. Active use of the metapleural glands by ants in controlling fungal infection. Proceedings of the Royal Society of London B: Biological Sciences 273: 1689–1695.

Halawani O, Dunn RR, Grunden AM & Smith AA. 2020. Bacterial exposure leads to variable mortality but not a measurable increase in surface antimicrobials across ant species. PeerJ 8: e10412.

Henrik H, Boomsma JJ & Tunlid A. 2014. Symbiotic adaptations in the fungal cultivar of leaf-cutting ants. Nature Communications 5: 5675.

Hoggard SJ, Wilson PD, Beattie AJ & Stow AJ. 2011. Social complexity and nesting habits are factors in the evolution of antimicrobial defences in wasps. PLOS ONE 6: e21763.

Jouvenaz DP, Blum MS & MacConnell JG. 1972. Antibacterial activity of venom alkaloids from the imported fire ant, Solenopsis invicta Buren. Antimicrobial Agents and Chemotherapy 2: 291–293.

Kappeler PM, Cremer S & Nunn CL. 2015. Sociality and health: Impacts of sociality on disease susceptibility and transmission in animal and human societies. Philisophical Transactions of the Royal Society B 370: 20140116.

Kesäniemi J, Koskimäki JJ & Jurvansuu J. 2019. Corpse management of the invasive Argentine ant inhibits growth of pathogenic fungi. Scientific Reports 9: 7593.

Koidsumi K. 1957. Antifungal action of cuticular lipids in insects. Journal of Insect Physiology 1: 40–51.

Lenth RV. 2025. emmeans: Estimated Marginal Means, aka Least-Squares Means.

Little AE, Murakami T, Mueller UG & Currie CR. 2006. Defending against parasites: Fungus-growing ants combine specialized behaviours and microbial symbionts to protect their fungus gardens. Biology Letters 2: 12–16.

Lockhart SR. 2019. Candida auris and multidrug resistance: Defining the new normal. Fungal Genetics and Biology 131: 103243.

MacGown JA. 2020. Mississippi Entomological Museum, Ants (Formicidae) of the Southeastern United States.

Marlier JF, Quinet Y & de Biseau JC. 2004. Defensive behaviour and biological activities of the abdominal secretion in the ant Crematogaster scutellaris (Hymenoptera: Myrmicinae). Behavioural Processes 67: 427–440.

Maschwitz U, Koob K & Schildknecht H. 1970. Ein beitrag zur funktion der metathoracaldrüse der ameisen. Journal of Insect Physiology 16: 387–404.

Maxson T & Mitchell DA. 2016. Targeted treatment for bacterial infections: Prospects for pathogen-specific antibiotics coupled with rapid diagnostics. Tetrahedron 72: 3609–3624.

Mendes TD, Borges WS, Rodrigues A, Solomon SE, Vieira PC, Duarte MCT & Pagnocca FC. 2013. Anti-Candida properties of urauchimycins from Actinobacteria associated with Trachymyrmex ants. BioMed Research International 2013: 835081.

Meunier J. 2015. Social immunity and the evolution of group living in insects. Philosophical Transactions of the Royal Society B: Biological Sciences 370: 20140102.

Moreau SJ. 2013. “It stings a bit but it cleans well”: Venoms of Hymenoptera and their antimicrobial potential. Journal of Insect Physiology 59: 186–204.

Penick CA, Halawani O, Pearson B, Mathews S, López-Uribe MM, Dunn RR & Smith AA. 2018. External immunity in ant societies: Sociality and colony size do not predict investment in antimicrobials. Royal Society Open Science 5: 171332.

Pie MR, Rosengaus RB & Traniello JF. 2004. Nest architecture, activity pattern, worker density and the dynamics of disease transmission in social insects. Journal of Theoretical Biology 226: 45–51.

Poulsen M, Bot AN, Nielsen MG & Boomsma JJ. 2002. Experimental evidence for the costs and hygienic significance of the antibiotic metapleural gland secretion in leaf-cutting ants. Behavioral Ecology and Sociobiology 52: 151–157.

Quinet Y, Vieira R, Sousa M, Evangelista-Barreto N, Carvalho F, Guedes M, Alves C, de Biseau J & Heredia A. 2012. Antibacterial properties of contact defensive secretions in neotropical Crematogaster ants. Journal of Venomous Animals and Toxins including Tropical Diseases 18: 441–445.

R Core Team. 2025. R: A Language and Environment for Statistical Computing.

Silverman J & Brightwell RJ. 2008. The Argentine ant: Challenges in managing an invasive unicolonial pest. Annual Review of Entomology 53: 231–252.

Singh N & Yeh PJ. 2017. Suppressive drug combinations and their potential to combat antibiotic resistance. The Journal of Antibiotics 70: 1033–1042.

Sosiak CE & Barden P. 2021. Multidimensional trait morphology predicts ecology across ant lineages. Functional Ecology 35: 139–152.

Stow A, Briscoe D, Gillings M, Holley M, Smith S, Leys R, Silberbauer T, Turnbull C & Beattie A. 2007. Antimicrobial defences increase with sociality in bees. Biology Letters 3: 422–424.

Téné N, Bonnafé E, Berger F, Rifflet A, Guilhaudis L, Ségalas-Milazzo I, Pipy B, Coste A, Leprince J & Treilhou M. 2016. Biochemical and biophysical combined study of bicarinalin, an ant venom antimicrobial peptide. Peptides 79: 103–113.

Touchard A, Aili SR, Fox EGP, Escoubas P, Orivel J, Nicholson GM & Dejean A. 2016. The Biochemical Toxin Arsenal from Ant Venoms. Toxins 8: 30.

Tragust S, Mitteregger B, Barone V, Konrad M, Ugelvig LV & Cremer S. 2013. Ants disinfect fungus-exposed brood by oral uptake and spread of their poison. Current Biology 23: 76–82.

Tragust S. 2016. External immune defence in ant societies (Hymenoptera: Formicidae): The role of antimicrobial venom and metapleural gland secretion. Myrmecological News 23: 119–128.

Vander Meer R. 2012. Ant interactions with soil organisms and associated semiochemicals. Journal of Chemical Ecology 38: 728–745.

Wang L, Elliott B, Jin X, Zeng L & Chen J. 2015. Antimicrobial properties of nest volatiles in red imported fire ants, Solenopsis invicta (Hymenoptera: Formicidae). The Science of Nature 102: 66.

Woolhouse M, Ward M, van Bunnik B & Farrar J. 2015. Antimicrobial resistance in humans, livestock and the wider environment. Philosophical Transactions of the Royal Society B: Biological Sciences 370: 20140083.

Yacoub T, Rima M, Karam M, Sabatier JM & Fajloun Z. 2020. Antimicrobials from venomous animals: An overview. Molecules 25: 2402.

Yan Y, An Y, Wang X, Chen Y, Jacob MR, Tekwani BL, Dai L & Li XC. 2017. Synthesis and Antimicrobial Evaluation of Fire Ant Venom Alkaloid Based 2-Methyl-6-alkyl-Δ1,6-piperideines. Journal of Natural Products 80: 2795–2798.

Yek SH, Nash DR, Jensen AB & Boomsma JJ. 2012. Regulation and specificity of antifungal metapleural gland secretion in leaf-cutting ants. Proceedings of the Royal Society of London B: Biological Sciences: rspb20121458.

Zeng Y, Hu XP & Suh SJ. 2016. Characterization of antibacterial activities of eastern subterranean termite, Reticulitermes flavipes, against human pathogens. PLOS ONE 11: e0162249.

