## Supplementary material for "Dual strategies in ant social immunity: Chemical diversity and pathogen specificity in antimicrobial defense": Table S1

### Supporting Information

**Table S1.** Colony collection information

| Ant species | Colony ID | Colony Location | Latitude | Longitude |
|---|---|---|---|---|
| *Brachyponera chinensis* | CBC0002 | Georgia | 34.062 | -84.6031 |
| | CBC0003 | Georgia | 34.0612 | -84.6031 |
| | CBC0004 | Georgia | 34.0354 | -84.5851 |
| | CBC0005 | Georgia | 34.0359 | -84.5863 |
| | CBC0006 | Georgia | 34.0368 | -84.5867 |
| | CBC0007 | Georgia | 34.0624 | -84.6034 |
| | CBC0008 | Georgia | 34.0347 | -84.5846 |
| | CBC0009 | Georgia | 34.0353 | -84.5845 |
| | CBC0010 | Georgia | 33.5242 | -84.3323 |
| | CBC0011 | Georgia | 33.5251 | -84.3306 |
| | CBC0014 | Georgia | 33.5219 | -84.3297 |
| | CBC0015 | Georgia | 33.5242 | -84.3323 |
| | CBC0016 | Georgia | 33.525 | -84.3303 |
| | CBC0017 | Georgia | 33.5219 | -84.3297 |
| *Crematogaster ashmeadi* | CCA0001 | Georgia | 34.0137 | -84.0855 |
| | CCA0002 | Georgia | 34.0613 | -84.6034 |
| | CCA0003 | Georgia | 34.0613 | -84.6034 |
| | CCA0004 | Georgia | 34.0612 | -84.6038 |
| | CCA0007 | Georgia | 34.0611 | -84.604 |
| | CCA0008 | Georgia | 34.0612 | -84.6042 |
| | CCA0009 | Georgia | 34.0615 | -84.6045 |
| | CCA0010 | Georgia | 34.0622 | -84.6039 |
| | CCA0011 | Georgia | 34.061 | -84.6037 |
| | CCA0012 | Georgia | 33.4418 | -84.1217 |
| | CCA0013 | Georgia | 33.4418 | -84.1217 |
| | CCA0014 | Georgia | 33.9712 | -84.1871 |
| | CCA0015 | Georgia | 33.9712 | -84.1854 |
| | CCA0016 | Georgia | 34.0357 | -84.585 |
| | CCA0017 | Georgia | 34.0366 | -84.5851 |
| *Dorymyrmex bureni* | CDB0002 | Florida | 30.7004 | -87.1091 |
| | CDB0003 | Florida | 30.7025 | -87.1091 |
| | CDB0004 | Florida | 30.6982 | -87.1091 |
| | CDB0005 | Georgia | 34.0461 | -83.9482 |
| | CDB0006 | Georgia | 34.0467 | -83.9488 |
| | CDB0007 | Georgia | 34.047 | -83.9489 |
| | CDB0008 | Georgia | 32.4521 | -84.9831 |
| | CDB0009 | Georgia | 32.4512 | -84.9816 |
| | CDB0011 | Georgia | 32.4521 | -84.9831 |
| | CDB0012 | Georgia | 32.4512 | -84.9816 |
| | CDB0013 | Georgia | 32.4521 | -84.9831 |

| | | | | |
|---|---|---|---|---|
| | CDB0014 | Georgia | 32.4521 | -84.9831 |
| | CDB0015 | Georgia | 32.4526 | -84.9881 |
| | CDB0016 | Georgia | 32.4539 | -84.9834 |
| | CDB0017 | Georgia | 32.454 | -84.9805 |
| | CDB0018 | Georgia | 32.4524 | -84.9823 |
| | CDB0019 | Georgia | 34.0482 | -83.9496 |
| *Linepithema humile* | CLH0001 | Georgia | 34.0354 | -84.5636 |
| | CLH0002 | Georgia | 34.0354 | -84.5835 |
| | CLH0003 | Georgia | 34.0614 | -84.6039 |
| | CLH0004 | Georgia | 34.0367 | -84.5838 |
| | CLH0005 | Georgia | 34.0373 | -84.5843 |
| | CLH0006 | Georgia | 33.5223 | -84.3128 |
| | CLH0007 | Georgia | 33.522 | -84.3134 |
| | CLH0008 | Alabama | 32.6087 | -85.4512 |
| | CLH0009 | Alabama | 32.6086 | -85.4511 |
| | CLH0010 | Georgia | 34.0017 | -84.5008 |
| | CLH0011 | Georgia | 34.0014 | -84.502 |
| | CLH0012 | Georgia | 34.0016 | -84.5017 |
| | CLH0013 | Georgia | 34.002 | -84.5017 |
| | CLH0014 | Georgia | 34.0372 | -84.5838 |
| | CLH0015 | Georgia | 34.0392 | -84.5829 |
| | CLH0016 | Georgia | 33.442 | -84.1214 |
| | CLH0017 | Georgia | 34.0014 | -84.502 |
| *Prenolepis imparis* | CPI0001 | Georgia | 34.0356 | -84.5853 |
| | CPI0002 | Georgia | 33.2463 | -83.9255 |
| | CPI0003 | Georgia | 33.2473 | -83.9267 |
| | CPI0004 | Georgia | 34.0357 | -84.5851 |
| | CPI0005 | Georgia | 34.0351 | -84.5854 |
| | CPI0006 | Georgia | 34.0356 | -84.5851 |
| | CPI0007 | Georgia | 34.036 | -84.5853 |
| | CPI0008 | Alabama | 32.5755 | -85.9592 |
| | CPI0009 | Alabama | 32.5763 | -85.9587 |
| | CPI0010 | Georgia | 33.248 | -83.9205 |
| | CPI0011 | Georgia | 34.0367 | -84.5851 |
| | CPI0012 | Georgia | 34.0348 | -84.5856 |
| | CPI0013 | Georgia | 33.443 | -84.1215 |
| | CPI0014 | Georgia | 33.4448 | -84.1199 |
| | CPI0015 | Georgia | 33.5671 | -84.1456 |
| | CPI0016 | Georgia | 33.5663 | -84.1453 |
| *Solenopsis invicta* | CSI0001 | Georgia | 34.0607 | -84.5916 |
| | CSI0002 | Georgia | 33.4993 | -84.2481 |
| | CSI0003 | Georgia | 33.5246 | -84.3298 |
| | CSI0004 | Georgia | 33.5673 | -84.1428 |
| | CSI0005 | Georgia | 34.0367 | -84.5838 |

| | | | | |
|---|---|---|---|---|
| | CSI0006 | Georgia | 34.033 | -84.5741 |
| | CSI0007 | Georgia | 34.0358 | -84.5836 |
| | CSI0008 | Georgia | 34.0332 | -84.5821 |
| | CSI0009 | Georgia | 34.0334 | -84.5823 |
| | CSI0010 | Georgia | 33.4928 | -84.2376 |
| | CSI0011 | Georgia | 34.0367 | -84.5838 |
| | CSI0012 | Georgia | 34.0367 | -84.5838 |
| | CSI0013 | Georgia | 34.0615 | -84.6034 |
| | CSI0014 | Georgia | 34.0625 | -84.6038 |
| | CSI0016 | Georgia | 34.0291 | -84.5716 |
| | CSI0019 | Georgia | 34.0337 | -84.5747 |
| | CSI0020 | Georgia | 34.0329 | -84.5746 |
| | CSI0021 | Georgia | 34.0328 | -84.5738 |
| | CSI0022 | Georgia | 34.0315 | -84.5731 |
| | CSI0023 | Georgia | 33.5223 | -84.3155 |
| | CSI0024 | Georgia | 33.5222 | -84.3127 |
| | CSI0025 | Georgia | 33.5235 | -84.3262 |
| | CSI0026 | Georgia | 33.5247 | -84.3288 |
| | CSI0027 | Georgia | 33.5245 | -84.3302 |
| | CSI0028 | Georgia | 33.5235 | -84.3293 |
| | CSI0029 | Georgia | 33.5232 | -84.3306 |
| | CSI0030 | Georgia | 33.5232 | -84.3087 |
| | CSI0031 | Georgia | 33.526 | -84.3102 |
| | CSI0032 | Georgia | 34.0332 | -84.5822 |
| | CSI0033 | Georgia | 33.5242 | -84.3156 |
| | CSI0034 | Georgia | 33.5252 | -84.3141 |
| | CSI0035 | Georgia | 33.4803 | -84.2265 |
| | CSI0036 | Georgia | 33.4791 | -84.2272 |
| | CSI0037 | Georgia | 33.48 | -84.2284 |
| | CSI0038 | Georgia | 33.4817 | -84.229 |
| | CSI0039 | Georgia | 33.4421 | -84.1214 |
| | CSI0040 | Georgia | 33.4425 | -84.1214 |
| | CSI0041 | Georgia | 33.567 | -84.1451 |
| | CSI0042 | Georgia | 32.451 | -84.9814 |
| | CSI0043 | Georgia | 32.4501 | -84.9818 |
| | CSI0044 | Georgia | 32.45 | -84.983 |
| | CSI0045 | Georgia | 32.4511 | -84.984 |
| | CSI0046 | Georgia | 34.0342 | -84.5827 |

**Table S2.** Sample sizes for antimicrobial assay for each test microbe and solvent combination (EtOH=ethanol; Iso=isopropanol; DCM=dichloromethane; Hex=hexane) showing number of independent nests per comparison.

| | *Staphylococcus epidermidis* | | | | *Escherichia coli* | | | | *Candida auris* | | | |
|---|---|---|---|---|---|---|---|---|---|---|---|---|
| **Ant species** | EtOH | Iso | DCM | Hex | EtOH | Iso | DCM | Hex | EtOH | Iso | DCM | Hex |
| *Brachyponera chinensis* | 14 | 14 | 14 | 14 | 14 | 14 | 14 | 14 | 14 | 14 | 14 | 14 |
| *Crematogaster ashmeadi* | 15 | 15 | 14 | 15 | 12 | 12 | 11 | 12 | 11 | 11 | 11 | 11 |
| *Dorymyrmex bureni* | 17 | 17 | 11 | 13 | 17 | 17 | 11 | 13 | 17 | 17 | 11 | 13 |
| *Linepithema humile* | 14 | 14 | 15 | 12 | 12 | 12 | 15 | 12 | 10 | 10 | 15 | 12 |
| *Prenolepis imparis* | 16 | 16 | 16 | 16 | 16 | 16 | 16 | 16 | 15 | 15 | 15 | 15 |
| *Solenopsis invicta* | 43 | 43 | 36 | 38 | 38 | 38 | 35 | 36 | 32 | 32 | 29 | 30 |

**Table S3.** Percent inhibition for ant extracts against each test microbe and solvent (EtOH=ethanol; Iso=isopropanol; DCM=dichloromethane; Hex=hexane).

| | *Staphylococcus epidermidis* | | | | *Escherichia coli* | | | | *Candida auris* | | | |
|---|---|---|---|---|---|---|---|---|---|---|---|---|
| **Ant species** | EtOH | Iso | DCM | Hex | EtOH | Iso | DCM | Hex | EtOH | Iso | DCM | Hex |
| *Brachyponera chinensis* | 14 | 0 | 0 | 0 | 43 | 57 | 0 | 0 | 79 | 79 | 7 | 0 |
| *Crematogaster ashmeadi* | 87 | 80 | 43 | 47 | 75 | 92 | 55 | 42 | 100 | 82 | 45 | 73 |
| *Dorymyrmex bureni* | 59 | 53 | 82 | 77 | 59 | 53 | 82 | 85 | 59 | 53 | 64 | 77 |
| *Linepithema humile* | 0 | 0 | 0 | 0 | 0 | 0 | 0 | 0 | 0 | 0 | 0 | 0 |
| *Prenolepis imparis* | 81 | 81 | 25 | 56 | 50 | 50 | 44 | 56 | 80 | 80 | 47 | 60 |
| *Solenopsis invicta* | 91 | 93 | 31 | 29 | 82 | 89 | 26 | 22 | 88 | 97 | 34 | 37 |

**Table S4.** Model results for pairwise comparisons of solvent within each species, with BH correction.

| Ant species | Comparison | Estimate | SE | Z-ratio | P-value |
|---|---|---|---|---|---|
| *Brachyponera chinensis* | DCM - EtOH | -4.18 | 1.12 | -3.729 | 0.0006 *** |
| | DCM - Hexane | 14.6 | 1480 | 0.01 | 1.0000 |
| | DCM - Iso | -4.18 | 1.12 | -3.729 | 0.0006 *** |
| | EtOH - Hexane | 18.8 | 1480 | 0.013 | 1.0000 |
| | EtOH - Iso | 0 | 0.546 | 0 | 1.0000 |
| | Hexane - Iso | -18.8 | 1480 | -0.013 | 1.0000 |
| *Crematogaster ashmeadi* | DCM - EtOH | -2.03 | 0.589 | -3.45 | 0.0034 ** |
| | DCM - Hexane | -0.229 | 0.47 | -0.488 | 0.7438 |
| | DCM - Iso | -1.82 | 0.561 | -3.241 | 0.0036 ** |
| | EtOH - Hexane | 1.8 | 0.583 | 3.092 | 0.0040 ** |
| | EtOH - Iso | 0.215 | 0.657 | 0.327 | 0.7438 |
| | Hexane - Iso | -1.59 | 0.555 | -2.864 | 0.0063 ** |
| *Dorymyrmex bureni* | DCM - EtOH | 0.784 | 0.496 | 1.58 | 0.1712 |
| | DCM - Hexane | -0.216 | 0.568 | -0.379 | 0.7045 |
| | DCM - Iso | 1.02 | 0.494 | 2.072 | 0.0814 • |
| | EtOH - Hexane | -1 | 0.489 | -2.047 | 0.0814 • |
| | EtOH - Iso | 0.239 | 0.4 | 0.599 | 0.6593 |
| | Hexane - Iso | 1.24 | 0.486 | 2.549 | 0.0649 • |
| *Linepithema humile* | DCM - EtOH | 0.0602 | 14600 | 0 | 1.0000 |
| | DCM - Hexane | 0.836 | 18800 | 0 | 1.0000 |
| | DCM - Iso | 0.241 | 15400 | 0 | 1.0000 |
| | EtOH - Hexane | 0.775 | 19600 | 0 | 1.0000 |
| | EtOH - Iso | 0.181 | 16400 | 0 | 1.0000 |
| | Hexane - Iso | -0.594 | 20200 | 0 | 1.0000 |
| *Prenolepis imparis* | DCM - EtOH | -1.36 | 0.444 | -3.077 | 0.0063 ** |
| | DCM - Hexane | -0.795 | 0.426 | -1.867 | 0.1239 |
| | DCM - Iso | -1.36 | 0.444 | -3.077 | 0.0063 ** |
| | EtOH - Hexane | 0.57 | 0.439 | 1.297 | 0.2337 |
| | EtOH - Iso | 8.00E-06 | 0.456 | 0 | 1.0000 |
| | Hexane - Iso | -0.57 | 0.439 | -1.297 | 0.2337 |
| *Solenopsis invicta* | DCM - EtOH | -2.76 | 0.356 | -7.737 | 0.0001 *** |
| | DCM - Hexane | 0.0575 | 0.309 | 0.186 | 0.8525 |
| | DCM - Iso | -3.46 | 0.43 | -8.043 | 0.0001 *** |
| | EtOH - Hexane | 2.81 | 0.355 | 7.921 | 0.0001 *** |
| | EtOH - Iso | -0.702 | 0.461 | -1.522 | 0.1536 |
| | Hexane - Iso | -3.52 | 0.429 | -8.192 | 0.0001 *** |

**Table S5.** Model results for pairwise comparisons of microbe within each species, with BH correction.

| Ant species | Comparison | Estimate | SE | Z-ratio | P-value |
|---|---|---|---|---|---|
| *Brachyponera chinensis* | *C. auris - E. coli* | 1.3762 | 0.584 | 2.356 | 0.0185 * |
| | *C. auris - S. epidermidis* | 3.9205 | 0.863 | 4.541 | 0.0001 *** |
| | *E. coli - S. epidermidis* | 2.5443 | 0.823 | 3.092 | 0.0030 ** |
| *Crematogaster ashmeadi* | *C. auris - E. coli* | 0.533 | 0.502 | 1.061 | 0.4328 |
| | *C. auris - S. epidermidis* | 0.6093 | 0.478 | 1.275 | 0.4328 |
| | *E. coli - S. epidermidis* | 0.0764 | 0.448 | 0.17 | 0.8647 |
| *Dorymyrmex bureni* | *C. auris - E. coli* | -0.2397 | 0.4 | -0.599 | 0.8400 |
| | *C. auris - S. epidermidis* | -0.1581 | 0.398 | -0.397 | 0.8400 |
| | *E. coli - S. epidermidis* | 0.0816 | 0.404 | 0.202 | 0.8400 |
| *Linepithema humile* | *C. auris - E. coli* | 1.0816 | 17000 | 0 | 1.0000 |
| | *C. auris - S. epidermidis* | 0.1999 | 12700 | 0 | 1.0000 |
| | *E. coli - S. epidermidis* | -0.8818 | 17100 | 0 | 1.0000 |
| *Prenolepis imparis* | *C. auris - E. coli* | 0.7476 | 0.386 | 1.937 | 0.1583 |
| | *C. auris - S. epidermidis* | 0.2679 | 0.39 | 0.688 | 0.4916 |
| | *E. coli - S. epidermidis* | -0.4797 | 0.372 | -1.289 | 0.2962 |
| *Solenopsis invicta* | *C. auris - E. coli* | 0.6234 | 0.324 | 1.926 | 0.1622 |
| | *C. auris - S. epidermidis* | 0.2013 | 0.315 | 0.638 | 0.5233 |
| | *E. coli - S. epidermidis* | -0.4222 | 0.303 | -1.395 | 0.2445 |

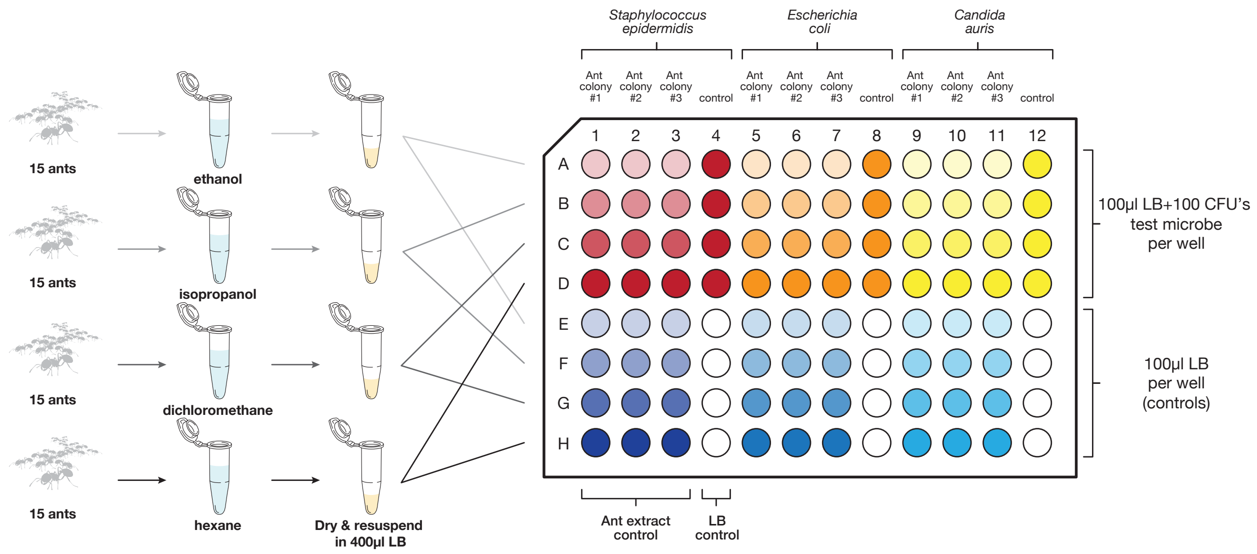


**Figure S1.** Plate diagram for antimicrobial assay. We soaked 15 ants of each species in one of four solvents. After drying, the solvents were resuspended in 400 µL of LB. Each extract was split evenly between a test well and a control well. Test wells contained 100 µL of LB with 100 CFUs of the respective test microbe, while control wells contained 100 µL of LB alone. Each plate included positive growth controls (microbes without ant extract) and LB-only controls to confirm the absence of contamination.
